# High-resolution experimental and computational electrophysiology reveals weak β-lactam binding events in the porin PorB

**DOI:** 10.1101/303891

**Authors:** Annika Bartsch, Salomé Llabrés, Florian Pein, Christof Kattner, Markus Schön, Manuel Diehn, Mikio Tanabe, Axel Munk, Ulrich Zachariae, Claudia Steinem

## Abstract

The permeation of most antibiotics through the outer membrane of Gram-negative bacteria occurs through porin channels. To design drugs with increased activity against Gram-negative bacteria in the face of the antibiotic resistance crisis, the strict constraints on the physicochemical properties of the permeants imposed by these channels must be better understood. Here we show that a combination of high-resolution electrophysiology, new noise-filtering analysis protocols and atomistic biomolecular simulations reveals weak binding events between the β-lactam antibiotic ampicillin and the porin PorB from the pathogenic bacterium *Neisseria meningitidis*. In particular, an asymmetry often seen in the electrophysiological characteristics of ligand-bound channels is utilised to characterise the binding site and molecular interactions in detail, based on the principles of electro-osmotic flow through the channel. Our results provide a rationale for the determinants that govern the binding and permeation of zwitterionic antibiotics in anion-selective porin channels.

## INTRODUCTION

All biological cells are enclosed by lipid bilayers. The translocation of hydrophilic molecules, such as nutrients and metabolic waste products, across these membranes is often rate-limiting for cellular growth and survival. Highly specialized proteins form hydrophilic channels across lipid bilayers which facilitate the exchange of these molecules.^1^ In bacteria, membrane channels also enable the inward permeation of antibiotics, which are usually too hydrophilic to pass directly through the lipid bilayer.^2-4^ In particular, antibiotics with activity against Gram-negative bacteria must be capable of permeating porin channels, which are located in the outer membrane of their cell envelope.^5^ This requirement not only imposes tight constraints on the physicochemical properties of permeants, but also enables pathogenic bacteria to evolve mutated porins, incurring a substantial decrease or complete loss of antibiotic uptake rates.^6-8^

The current rise in antimicrobial resistance urges the development of drugs that are able to more readily permeate the outer membrane of Gram-negative bacteria,^9-12^ as these organisms constitute the vast majority of priority pathogens exhibiting multi-or pan-drug resistance.^13,14^ Yet, due to the intrinsically low permeability of most antibiotics across the outer membrane and its channel proteins, too few newly developed drug candidates show activity against Gram-negative bacteria.^15,16^ It is therefore essential to better understand the physicochemical restraints governing antibiotic permeation and binding in porins in order to design drugs with improved permeability.^17-19^ At the same time, this insight will also help to elucidate how channel mutations decrease antibiotic uptake by altering drug-channel interactions and thereby contribute to the development of resistance.^9,20^ The decreased influx rates of antibiotics into the bacteria caused by altered antibiotic-channel interactions can be aggravated by the overexpression of efflux pumps, which in combination, often leads to highly resistant phenotypes^21^. In order to aim for lower-barrier, rapid inward translocation rates, it is especially important to determine whether binding in the channel occurs. However, interactions between drugs and porins are often weak and short-lived, and therefore difficult to characterise.

In this study, we demonstrate that a combination of high-resolution single-channel electrophysiology, novel high-sensitivity electrophysiology analysis methods,^22^ and atomistic molecular dynamics simulations allowed us to reveal in structural detail a drug-channel binding interaction that is too weak to be captured by X-ray crystallography. We set out to investigate the interaction between ampicillin and PorB from *Neisseria meningitidis*, the pathogen that causes bacterial meningitis. PorB is the second most abundant protein in its outer membrane and a key component in both its pathogenicity and the activation of the host immune response.^23-25^ Moreover, PorB plays an essential role in the drug-resistance of *Neisseria* since it is the main inward route for antibiotics,^26^ and mutations in the pore-lining residues are linked to resistance to β-lactamic antibiotics.^8,27^ By contrast to other well-known general porins from Gram-negative bacteria such as OmpF from *E. coli*, which are strongly cation-selective, PorB belongs to a small set of porins showing strong selectivity for anions^23,28,29^. Therefore, although the binding mode of a β-lactam to the general porin OmpF from *E. coli* has previously been elucidated^30^, differences in the interactions between PorB and β-lactamic antibiotics are likely.

A variety of techniques are used to study the interplay between small-molecules and membrane channels.^5^ X-ray crystallography provides structural insight into high-affinity molecular binding interactions. Furthermore, liposome-swelling assays qualitatively measure influx rates of small-molecules through channel proteins. Electrophysiological measurements are capable of quantifying interactions between porin channels and antibiotics by revealing channel current disruptions caused by drug interactions with the pore. However, current fluctuations induced by small molecules can either reflect their low-affinity binding and/or their transient passage through the channel.^4,31^ In order to provide structural information at sufficient detail for improved drug design,^19,32^ it is important to reliably identify these scenarios, and to characterise the molecular determinants of any potential binding event between the antibiotic molecule and the protein channel.^33,34^ Here we demonstrate how we have combined new and traditional methodologies of membrane channel characterisation to achieve this.

## RESULTS AND DISCUSSION

First, we performed single channel measurements of PorB at voltages between 40-120 mV with and without ampicillin. Since PorB is a trimeric porin, it exhibits three conductance states in the absence of ampicillin, which reflect the open state of a monomer (*G*_M_ = 0.44 ± 0.18 nS), dimer (*G*_D_ = 0.89 ± 0.26 nS) and trimer (*G*_t_ = 1.52 ± 0.46 nS) (**Fig. S1** and **S2**). The determined PorB trimer conductance *G*_T_ is similar to previously reported values of 1.0-1.5 nS.^35,36^ Furthermore, the sequential opening and closing of the channel observed here is in agreement with cooperative gating, as previously described in the literature.^36^

The open channel current reveals the interaction of the antibiotic with the pore by displaying short interruptions, which are only recorded in the presence of ampicillin (Figs. 1A/B). As a result, we observe an additional conductance state (Figs. 1C/D). The difference between the ampicillin-induced and the open channel sub-conductance states is defined as the “blocked” amplitude *G*_B_, which quantifies the extent by which the conductance of the open channel is reduced by the interaction with ampicillin. This “blocked” amplitude arises from a transient molecular binding event, which can result in translocation of the drug.^4,31^ We report a *G*_B_ = 1.18 ± 0.06 nS (Fig. 1E), which suggests a simultaneous interaction of ampicillin molecules with all three PorB monomers, as previously described, e.g. for chitoporin.^37^ The blocked amplitudes *G*_B_ display a slight dependence on the applied voltage but not on drug concentration (Fig. 1F).

**FIGURE 1.**
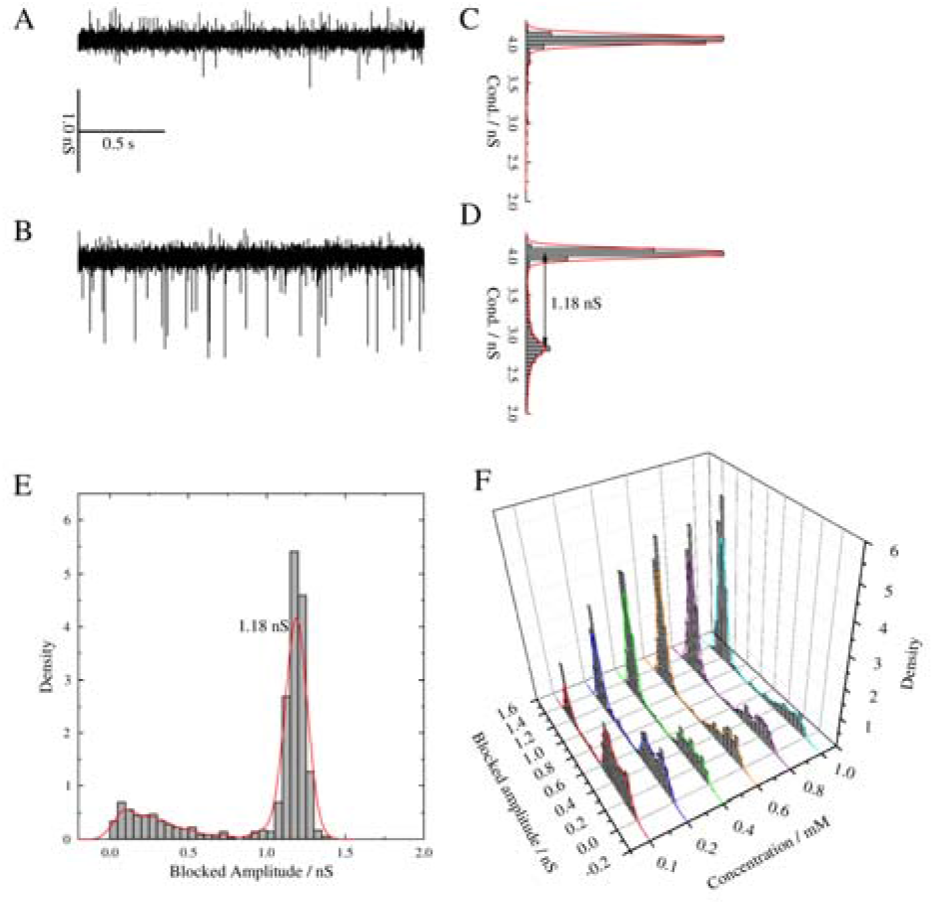
Representative conductance traces for PorB (**A**) before and (**B**) after addition of 1 mM ampicillin. The corresponding histograms (**C**) and (**D**) show the frequencies of the obtained conductance levels (Cond.). Recordings were obtained on BLMs (DPhPC/cholesterol, 9:1) in 1 M KCl, 10 mM HEPES, pH 7.5 with an applied voltage of +80 mV. (**E**) Event histogram of the amplitudes blocked by ampicillin with one maximum at G_B_ = 1.18 ± 0.06 nS, determined by a Gauss fit. The red line is the kernel density estimation using a Gaussian kernel. (**F**) Event histograms and kernel density estimation using a Gaussian kernel of the amplitudes blocked by ampicillin for PorB recorded at +80 mV. The ampicillin concentration was increased in a stepwise manner from 0.1-1.0 mM.

Since events induced by small molecules are short-lived in comparison to the open channel lifetime, high filter frequencies (10-15 kHz) are usually required to reduce the noise level of recorded signals and to improve the statistical analysis of the observed events.^4,26^ However, the filter length used by automated analysis methods limits the detection of short blockage events beyond the filter resolution, which means that relevant data is discarded as noise (missed events). Therefore, to be able to detect the short blockage events shown in Fig. 1B, we used a markedly reduced filtering frequency of 5 kHz and employed a novel model-free analysis method named JULES,^22^ which we have recently designed to robustly detect and reconstruct times and amplitudes of blockage events (**Figs. S3-S5**). JULES uses a combination of multi-resolution techniques and local deconvolution, allowing a precise idealization of events below the filter length; in particular amplitudes and residence times that are smoothed by the filter are reconstructed with high precision. Here, it enabled us to detect and idealise events down to a length of 80 μs. Based on this idealisation we were able to confirm a Markov model at this time resolution. Assuming now a Markov model on smaller time scales permits us to determine residence times down to about 20 μs using a missed event correction^22^. This improves previously achieved detection limits of about 50 μs^4,31,32^, which would not be sufficient for the subsequent analysis. In order to validate our analysis, we employed a hidden Markov model (HMM) in addition, in which the filter has been taken into account explicitly as well. In summary, our HMM analysis confirmed all findings by JULES; in particular, we found similar residence times. An analysis of even shorter time scales is often carried out by current distribution fitting (excess-noise analysis)^38,39^. However, these methods are based on the assumption of a HMM throughout the analysis and more importantly, their results are heavily influenced by model violations. A more detailed discussion and comparison of these methods can be found in the introduction of Pein et al. ^22^. Additionally, details of the models and the corresponding statistical analysis are given in the supporting information.

The voltage-dependence of the ampicillin interaction is confirmed by determining residence times at a fixed ampicillin concentration of 1 mM and a range of voltages between +(40-120) mV (Fig. 2A). These display a bell-shaped relationship with a maximum at around +100 mV. We further observe a linear decay of the frequency of blockage with increasing voltage (Fig. 2B).

**FIGURE 2.**
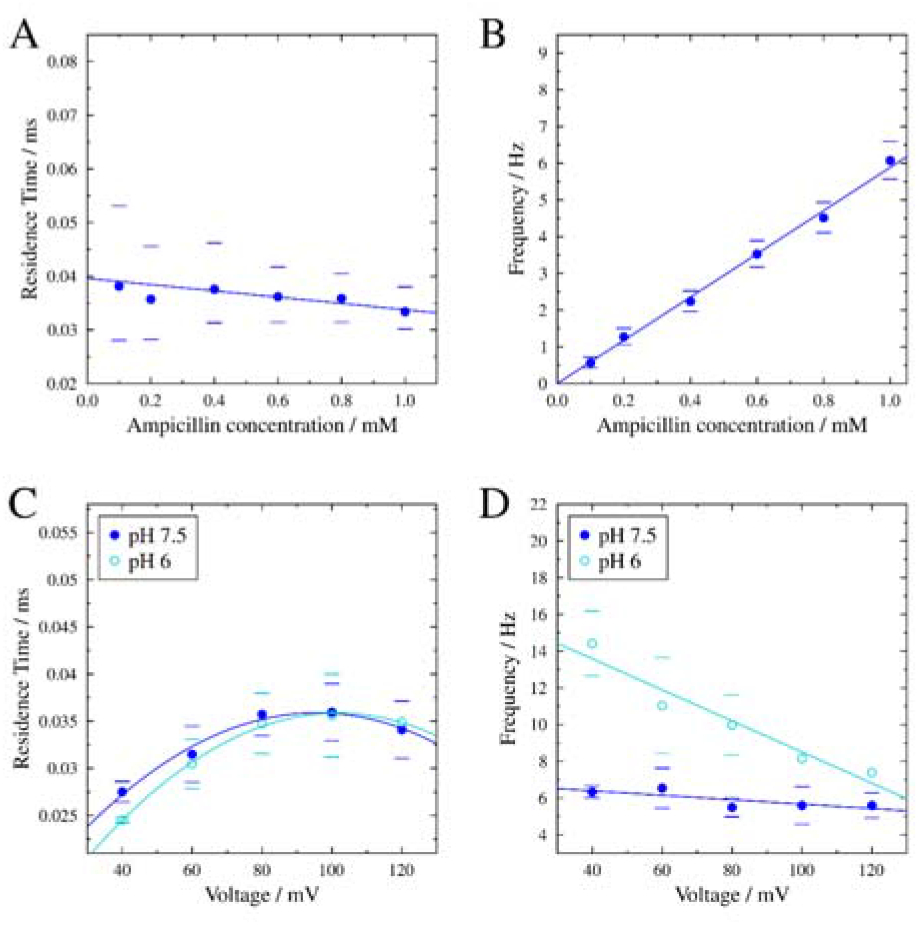
Characterisation of the residence times (**A, C**) and blockage frequencies (**B, D**) of ampicillin at different concentrations of ampicillin and different applied voltages. Parabolic and linear fits serve as a guide-to-the-eye. Averages of *n* = 4 (pH 7.5) and *n* = 2 (pH 6) measurements with the corresponding standard deviation are shown for the voltage dependence. Exemplar measurements including confidence intervals with 95% significance are depicted for the ligand concentration dependence.

We next investigated the kinetics of the interaction as a function of ampicillin concentration in a range of 0.1-1.0 mM and at a fixed voltage of +80 mV. The new analysis protocol described above enabled us to determine a residence time of τ_R_ = (35 ± 1) μs without any obvious dependence on ampicillin concentration (Fig. 2A). However, the frequency of blockage shows a linear dependence on the drug concentration (Fig. 2B), as expected for an increasing number of available antibiotic molecules. Therefore, a higher concentration of antibiotic enhances the probability of interaction. However, once an ampicillin molecule interacts with a PorB monomer, only an applied positive voltage of up to +100 mV can prolong the residence time of ampicillin. By contrast, the stabilizing effect observed at positive voltages is greatly attenuated at opposite membrane polarisation, up to the point that we observe an absence of blockage events at negative voltages.

Notably, even at high ampicillin concentrations, we observe a completely unilateral interaction with the channel. Blockage events are only detected when ampicillin is added to the same compartment as the protein, i.e., to the *cis* compartment. Since the extended extracellular loops of porins such as PorB are too polar to traverse the bilayer,^23^ PorB exclusively inserts into planar lipid bilayers in one direction,^40,41^ in which the extracellular site faces the side of insertion (*cis* side).

These findings therefore identify the sub-conductance state induced by the presence of ampicillin as a binding interaction,^31,42^ as opposed to a transient migration of the drug across the channel. Although a translocation process with largely unbalanced inward and outward rates cannot be completely discarded, the marked asymmetry of this interaction strongly suggests ampicillin binding only to PorB.^31^ Further, our electrophysiological data suggest that this binding site has the following characteristics: (1) it is readily accessible only from the extracellular compartment; (2) drug binding is able to obstruct the PorB pore, and (3) binding is stable only under positive voltages.

Having established a binding site for ampicillin in PorB, we applied molecular docking calculations followed by extended molecular dynamics simulations to achieve insight on this interaction at the atomistic level. PorB forms a 16-stranded β-barrel channel with short connecting turns on the periplasmic side and long loops on the extracellular side (crystal structure, PDB ID: 3VY8).^23,28^ The extracellular loop 3 (L3) folds back into the pore, constricting it to a diameter of 8 Å at its narrowest point (Fig. 3). Here, the acidic residues E116 and E110 and a cluster of conserved basic residues (K64, R77, K100 and R130) create a strong transverse electrostatic field, likely to interact favorably with zwitterionic molecules such as ampicillin (**Fig. S6**).^32,43^ Due to our electrophysiological findings, we focused on the extracellular side of the constriction zone as the most probable binding region for the drug. Our docking calculations identified a stable binding mode in this region, in which the zwitterionic form of ampicillin is located at the base of the extracellular vestibule of the channel (Fig. 3A) and largely obstructs the entrance to the constriction zone. Of note, the anionic state of ampicillin interacts with PorB at the same site and in the same pose (**Fig. S7**). Extended equilibrium molecular dynamics simulations further confirm the stability of this binding mode.

**FIGURE 3.**
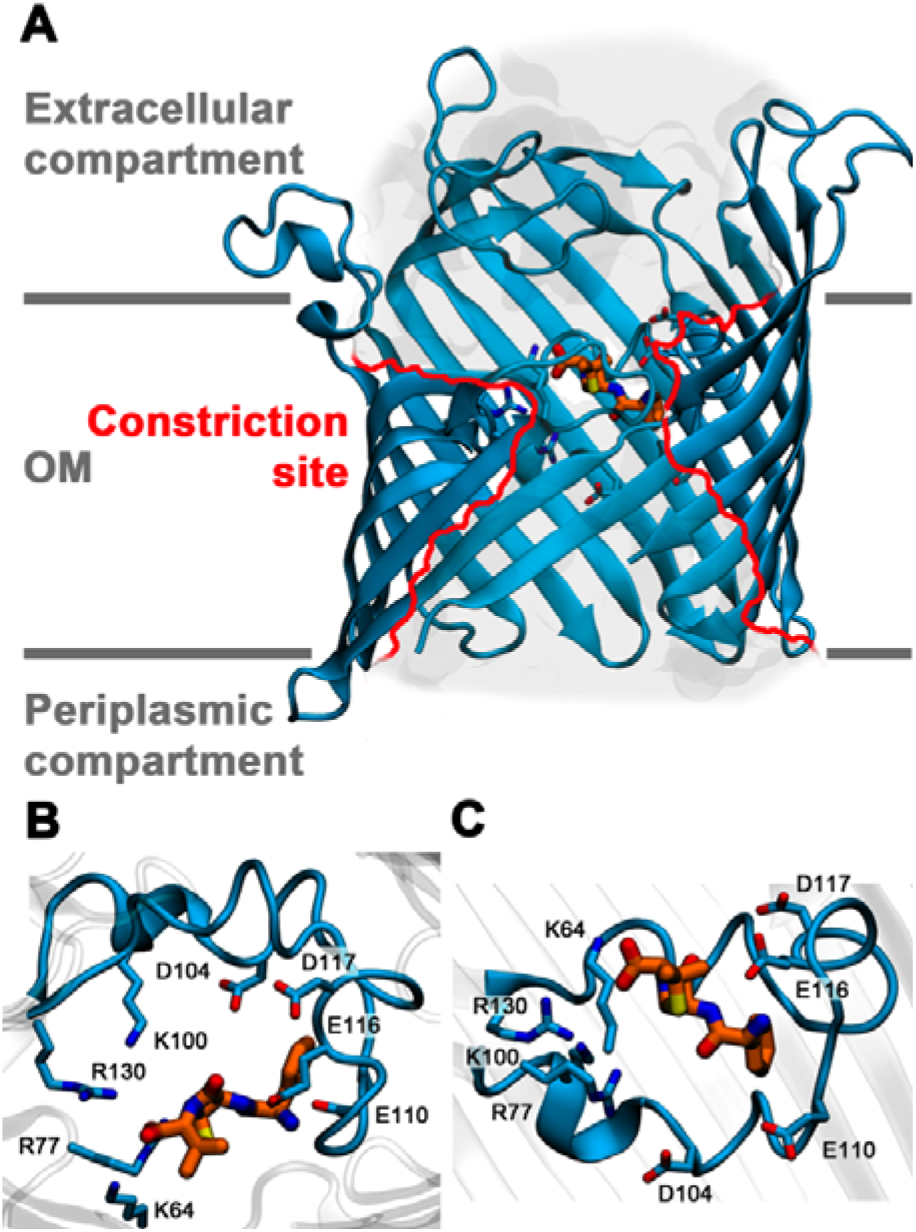
(**A**) Asymmetric location of the binding site on the extracellular side of the PorB eyelet. The constriction zone is highlighted by red lines. The protein and ampicillin carbon atoms are shown in blue cartoon and orange sticks, respectively. Close-up view of the binding mode of ampicillin seen from (**B**) the extracellular vestibule and (**C**) from the membrane. The L3 loop of PorB is shown in blue cartoon and blue sticks and the ampicillin carbon atoms are shown as orange sticks.

At pH 6, ampicillin is almost exclusively zwitterionic, while at pH 7.5, a mixture of the zwitterionic and anionic form is present (**Fig. S5**). Within the pore environment of PorB, we anticipate that the major species interacting with the channel is the zwitterionic form at both pH conditions. The protonation state is likely to be influenced by the microenvironment of the pore created by the transverse electrostatic field, and the favourable interactions between the small-molecule and the protein are substantially increased in its zwitterionic form (**Fig. S6**). As a result, the residence times should not be affected by the change in pH, which is in agreement with our experimental results shown in Fig. 2C. We find a difference in the frequency of blockage depending on the experimental conditions (Fig. 2D), which however can be attributed to factors beyond the actual binding site of ampicillin, e.g. elastic structural deformations caused by the applied electric field^44,45^.

In the docked configuration, the ampicillin molecule extends along the pore axis and interacts both with the acidic and basic clusters of the constriction zone. While the acid moiety of ampicillin forms a salt bridge with residues R77 and K64 on the basic side of the PorB eyelet (Fig. 3B), its amino group establishes a salt bridge with the side chain of residue E116 and additionally binds to the carbonyl group of the backbone of residue G112, both located on the acidic side of the eyelet. The phenyl moiety of ampicillin is wedged in by a shallow hydrophobic site, formed by the aliphatic portions of the E116 and E110 side chains and the backbone of residues W109, E110, S111, W115, G120 and E116.

To more closely reflect the experimental conditions, we next performed further molecular dynamics simulations of PorB channels inserted into POPC membranes under applied membrane voltages using the Computational Electrophysiology protocol.^46,47^ We determined the conductance of the channels for Na^+^ and Cl^-^ with and without bound ampicillin in the simulations, and tested its dependence on the polarity of the electric field. In order to achieve improved sampling, we extended the voltage range to values of ~125 - ~500 mV.

In the absence of the antibiotic, we observe a PorB trimer conductance of *G*_T,comp_ = 168 ± 0.17 nS at positive voltages and *G*_T,comp_ = 180 ± 0.14 nS at negative voltages. The values are equal within their error margins and in agreement with the experimental values reported here and in previous computational studies^28,46^ (Fig. 4A).

**FIGURE 4.**
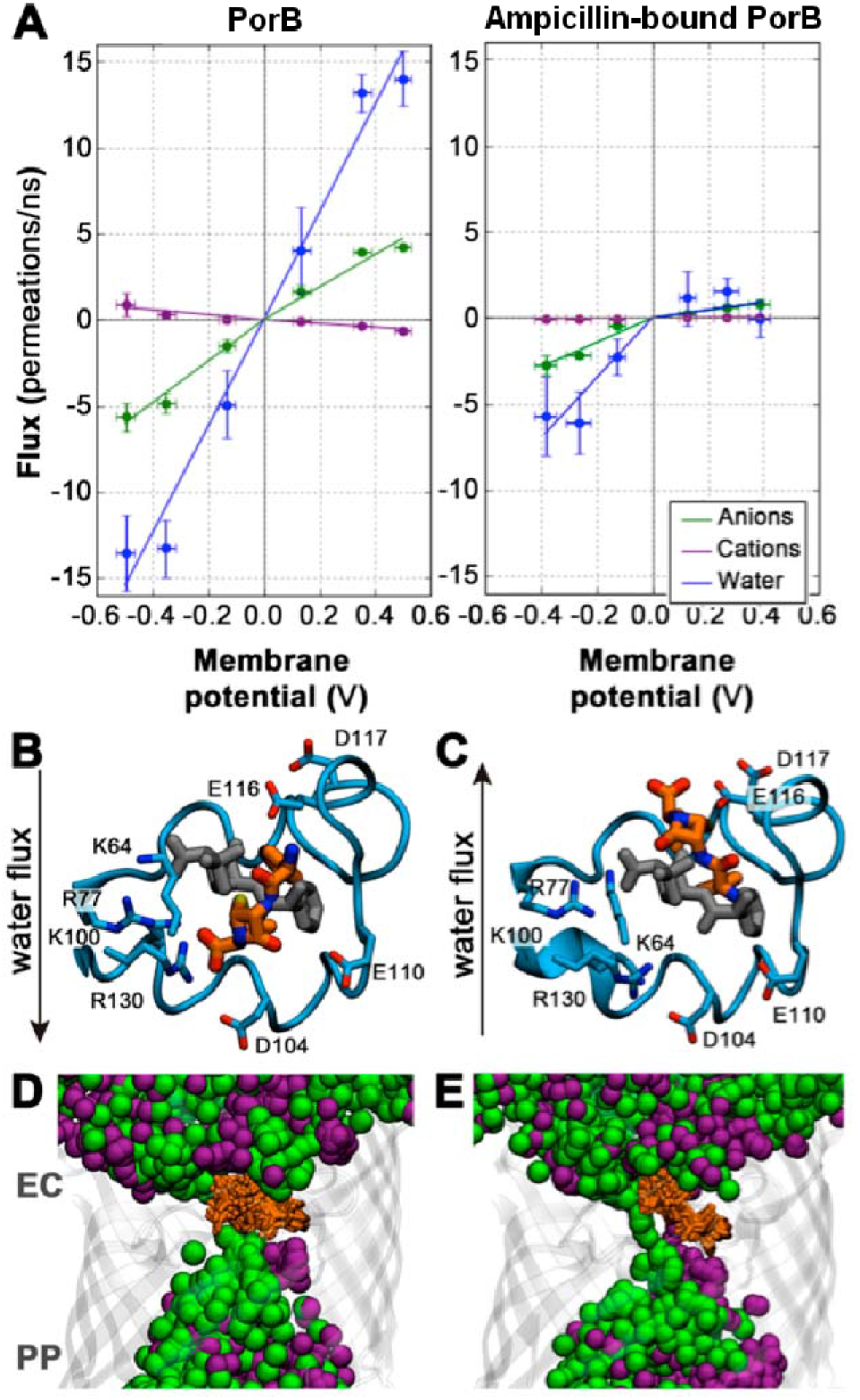
(**A**) Effect of ampicillin binding on the ion current in simulations under voltage. The asymmetric flux of Cl^-^ (green), Na^+^ ions (purple) and electro-osmotic flow of water molecules (blue) across the eyelet at negative and positive voltages is shown. Positive values of flux account for permeations from the extracellular compartment to the periplasmic compartment. Representative modes of ampicillin binding at positive (**B**) and negative voltages (**C**). The L3 loop of PorB is represented as blue cartoon and sticks and the initial and final structures of ampicillin are depicted as grey and orange sticks, respectively. (**D, E**) Overlay of ion positions (green spheres, Cl^-^; purple spheres, Na^+^) from 40 snapshots taken from the last 200 ns of simulation of bound ampicillin within PorB at (**D**) ~ +400 mV and (**E**) ~ −400 mV. Ion permeation is blocked at positive voltages, while anions can pass the PorB eyelet at negative voltages, leading to the asymmetric current-voltage behavior.

By stark contrast, presence of the drug abolishes the symmetry of the channel current-voltage relationship. When ampicillin resides at the identified binding site, the conductance of the complex at negative voltages is *G*_T-ampicilin,comp_ = 0.87 ± 0.28 nS, while that at positive voltages is reduced by ~67% to *G*_T-ampicillin,comp_ = 0.29 ± 0.10 nS. Importantly, at negative voltages, ampicillin is no longer stably locked in the binding site, which leads to a partial recovery of the conductance values of the apo channel (Fig. 4C), although it is not fully recovered on the limited time-scale of our simulations. These findings are in very good agreement with our experimental observations, since the location and topology of the ampicillin binding site results in asymmetric currents through PorB when the drug is bound in our simulations. The conductance at positive voltages shows a much more pronounced reduction due to drug binding than the conductance at negative voltages, owing to the destabilisation of the binding mode of ampicillin at negative voltages.

To elucidate the molecular basis for these effects, we investigated the details of ion flux across the channel. Since PorB is a strongly anion-selective channel, the bulk of the electric current arises from the flux of Cl’ ions, and a much smaller portion from the exchange of Na^+^ ions. The size of the constriction zone is wide enough to allow ions to traverse the pore in an at least partially hydrated state. During voltage-driven permeation, we find that Cl^-^ ions retain about half of their original hydration shell when they cross the constriction zone, such that 3.54 ± 0.16 water molecules are carried across the pore per Cl^-^ ion exchanged. The co-permeation of water, dragged through membrane channels by permeating ions, is known as the electro-osmotic flow (EOF).^33^

At positive voltages, the flux of water molecules is directed from the extracellular side of PorB toward its periplasmic face. In the case of ampicillin-bound PorB, the EOF stabilizes ampicillin binding since the water flux exerts forces on the antibiotic molecules driving them into the eyelet of PorB (Fig. 4B). In contrast, at negative voltages, the flow of water molecules is oriented from the periplasm towards the extracellular side. This inverted EOF destabilizes the binding of ampicillin to PorB by exerting a force on the molecule, which pushes the drug back into the extracellular lumen. In the simulations, this results in the loss of interactions between the acid moiety of ampicillin and the conserved basic cluster of the PorB eyelet (Fig. 4C). Ultimately, we expect the inverted EOF to completely remove ampicillin from its binding site and displace it into the extracellular compartment, although the full transition is not observed during the limited time-span of our simulations. The pronounced EOF effect provides further evidence that the binding mode of ampicillin to PorB is of relatively low affinity. The high level of agreement between our simulation results and our experimental findings suggests that we have correctly identified the site and mode of binding between the β-lactam antibiotic ampicillin and PorB at atomic level.

## CONCLUSION

In summary, we show here that a novel approach for analyzing high-resolution electrophysiological recordings, in combination with molecular docking calculations and molecular dynamics simulations under voltage, enables the detailed identification of a weak binding between a small-molecule antibiotic and a bacterial membrane channel. The new analysis method is able to correctly identify blockage events in data that would normally be discarded as noise. The efficiency of antibiotic binding or permeation across these channels determines the influx rate of the drugs into pathogenic bacteria and a reduction in inward permeation rates is often crucial for the development of antibiotic resistance. Therefore, the structural characterization of antibiotic-porin binding has important implications both for biomedicine and the drug design of new antibiotics. The binding site revealed by our approach is wedged between the clusters of conserved acidic and basic residues of the eyelet region, suggesting a common binding site for zwitterionic drugs within porin channels, in which ligand binding is strongly determined by the transverse electrostatic field.^32,43^ Importantly, we were able to use information on a marked asymmetry in the experimental currents across the drug-bound channel, which is often detected in ligand-bound membrane pores, and which has been shown to be caused by the electro-osmotic flow of water through the channel,^33,34,43,48^ to validate the binding mode and characterise the drug-channel interaction in detail. These results show that a combination of well-established techniques with new analysis protocols permits the identification and validation of previously undetectable low-affinity drug binding events in a biological channel protein. Furthermore, our findings shed light onto ampicillin binding in bacterial PorB, which is crucial to achieve a better understanding of the determinants of antibiotic permeation into bacteria and to overcome antibiotic resistant phenotypes that are linked to porin mutations.

## METHODS

### Electrophysiological setups and measurements

Single channel recordings of PorB were performed on solvent-free planar bilayers using the Port-a-Patch instrument (Nanion Technologies, Munich, Germany). Giant unilamellar vesicles (GUVs) composed of 1,2-diphytanoyl-*sn*-glycero-3-phosphocholine (DPhPC)/cholesterol (9:1) were prepared by electroformation (AC, *U* = 3 V, peak-to-peak, *f* = 5 Hz, *t* = 2 h) in the presence of 1 M sucrose at 20 °C.^49^ Spreading of a GUV in 1 M KCl, 10 mM HEPES, pH 7.5 on an aperture (*d* = 1-5 μm) in a borosilicate chip by applying 10-40 mbar negative pressure resulted in a solvent-free membrane with a resistance in the GΩ-range. Once the membrane with a GΩ-seal was formed, varying amounts of a PorB stock solution (2.2 μM in 200 mM NaCl, 20 mM Tris, 0.1 % (*w/w*) N,N-dimethyldodecylamine N-oxide (LDAO), pH 7.5) were added to the buffer solution (50 μL) at an applied DC potential of +40 mV. Current traces were recorded at a sampling rate of 10 kHz and filtered with a low-pass four-pole Bessel filter of 1 kHz using an Axopatch 200B amplifier (Axon Instruments, Union City, CA, USA). For digitalization, an A/D converter (Digidata 1322; Axon Instruments) was used and data analysis was performed with the Clampfit 10.4.0.36 from the pClamp 10 software package (Molecular Devices, Sunnyvale, CA, USA). Solvent-free bilayers were used to obtain the three conductance values of the PorB trimer (**Fig. S1**), as the PorB channel gates much more frequently than in black lipid membranes (BLMs) (**Fig. S2**). For electrophysiological measurements in the presence of ampicillin including the corresponding control experiments, BLMs were used. BLMs were prepared by adding 1-2 μL of lipids (DPhPC/cholesterol, 9:1) dissolved in n-decane (30 mg/mL) to an aperture (*d* = 50 μm) in a PTFE foil (DF100 cast film, Saint-Gobain Performance Plastics, Rochdale, UK) fixed between two cylindrical PTFE-chambers filled with 3.0 mL buffer (1 M KCl, 10 mM HEPES, pH 7.5 or pH 6). Protein was added to the *cis* chamber and inserted by stirring at an applied DC potential of +40 mV. After protein insertion, ampicillin was added from a stock solution (25 mM in 1 M KCl, 10 mM HEPES, pH 7.5 or pH 6.0) to both sides of the BLM. For control experiments, ampicillin was added only to the *trans* side. Current traces were recorded at a sampling rate of 50 kHz and filtered at 5 kHz.

### Analysis of current traces in the presence of ampicillin

Model-free idealizations are obtained by JULES.^22^ Its combination of multiresolution techniques and local deconvolution allows a precise idealisation of events below the filter length; in particular amplitudes and residence times that are smoothed by the filter are reconstructed with high precision (**Fig. S3**). In this manner, JULES extends the model-free idealization method JSMURF^50^ to scales below the filter length. The idealized blockage supports a two state Markov model. The conductance losses by blocking a single channel are determined by Gaussian fits of the amplitudes. Residence times and frequencies are determined by fitting the Markov model taking missed events into account. An analysis with a new Hidden Markov model approach, which is able to take the lowpass Bessel filter explicitly into account, confirms these results.

### Docking calculations

The binding mode of ampicillin was explored by means of docking calculations carried out with the GOLD^51^ and rDock^52,53^ software packages. The structural model of PorB considered in the docking calculations was the X-ray structure of PorB solved by Kattner et al. (PDB entry 3VY8).^28^ Both zwitterionic and anionic forms of ampicillin were subjected to 100 docking runs. Whereas the protein was kept rigid, GOLD and rDock account for the conformational flexibility of the ligand during docking. Resulting binding poses were analysed by visual inspection in conjunction with the docking scores.

### Simulation system set-up

PorB was modelled using the X-ray structure obtained by Kattner et al. (PDB entry 3VY8).^28^ PorB trimers were embedded into a preequilibrated 160×160 Å^2^ 1-palmitoyl-2-oleoyl-sn-glycero-3-phosphocholine (POPC) bilayer accounting for 942 POPC molecules. Water molecules forming a layer of 25 Å were added to each side of the bilayer, and Na^+^ and Cl^-^ were included to achieve an ionic strength of 1 M. The GROMACS utility membed^54,55^ was used to embed PorB trimers into a POPC bilayer. The aqueous solution consisted of about 57,000 water molecules and 1866 Na^+^ and 1896 Cl^-^ ions in both the apo and ampicillin-bound system. The Parm99SB force field,^56,57^ and virtual sites for hydrogen atoms^58^ were used for the protein. The POPC molecules were parameterized according to the lipid parameters derived by Berger et al.,^57,59^ the SPC/E water model was used to model water molecules^60^ and Joung and Cheatham parameters^61^ were used to model the ions. The zwitterionic ampicillin molecule was parameterized using the gaff force field^62^ in conjunction with RESP (HF/6-31G(d)) charges^63^ as implemented in the Antechamber module of the AMBER12 software package.^64^

### Molecular dynamics simulations

MD simulations were carried out with the GROMACS package, version 5.1.5.^65^ For each system, the geometry was minimised within four cycles that combined 3500 steps of the steepest descent algorithm followed by 4500 of conjugate gradient minimisation. Thermalisation of the system was performed in 6 steps of 5 ns; each step gradually increased the temperature from 50 K to 320 K, while the protein was restrained with a force constant of 10 kJ mol^-1^ Å^-2^. The systems were equilibrated for 100 ns keeping the protein restrained. Production runs consisted of trajectories of 200 ns length. The temperature was kept constant by weakly coupling (*t* = 0.1 ps) the membrane, protein, and solvent separately to a temperature bath of 320 K with the velocity-rescale thermostat of Bussi et al.^66^ The pressure was kept constant at 1 bar using semi-isotropic Berendsen coupling.^67^ Long-range electrostatic interactions were calculated using the smooth particle mesh Ewald method^68^ beyond a short-range Coulomb cut-off of 10 Å. A 10 Å cut-off was also used for Lennard-Jones interactions. The LINCS algorithm^69^ was used to restrain the system and the SETTLE algorithm^70^ was used to constrain bond lengths and angles of water molecules. Periodic boundary conditions were applied. The integration time-step was 4 fs, making use of the Berger lipid and the virtual sites models.

### Computational Electrophysiology simulations (CompEL)

Each system was duplicated along the z-axis to construct a double bilayer system, and ionic imbalances of 4, 8 and 12 Na^+^ ions were used between the aqueous compartments to generate a range of transmembrane potentials of ± 130, ± 350 and ± 500 mV, as previously described by Kutzner et al.^47^ Production runs consisted of trajectories of 200 ns length for the apo systems and 400ns for the ampicillin-bound systems. The applied membrane potential was calculated using the GROMACS utility gmx potential with overlapping 20-ns time windows.

### Analysis of the molecular dynamics trajectories

MDAnalysis^71^ and MDtraj^72^ were used to analyze root mean square deviation of the protein (RMSD), distances, flux of water molecules and computational conductance values.

## SUPPORTING MATERIAL

Including methodological detail and seven figures.

## AUTHOR CONTRIBUTIONS

A.B., C.K., M.S. and M.T performed the experiments, F.P., M.D. and A.M. developed and applied the mathematical tools to evaluate the data, S.L. performed and analyzed the MD simulations. S.L., U.Z. and C.S. wrote the manuscript with help from all other authors.

## ACKNOWLEDGEMENTS

We are grateful to N. Denkert and M. Meinecke for support in constructing and performing the measurements on black lipid membranes and I. Mey for helpful discussions. We acknowledge funding through the Wellcome Trust Interdisciplinary Research Funds (grant WT097818MF), the Scottish Universities’ Physics Alliance (SUPA) and the Tayside Charitable Trust.

